# Sensitivity enhancement and selection are shared mechanisms for spatial and feature-based attention

**DOI:** 10.1101/2021.01.26.428350

**Authors:** Daniel Birman, Justin L. Gardner

## Abstract

Human observers use cues to guide visual attention to the most behaviorally relevant parts of the visual world. Cues are often separated into two forms: those that rely on spatial location and those that use features, such as motion or color. These forms of cueing are known to rely on different populations of neurons. Despite these differences in neural implementation, attention may rely on shared computational principles, enhancing and selecting sensory representations in a similar manner for all types of cues. Here we examine whether evidence for shared computational mechanisms can be obtained from how attentional cues enhance performance in estimation tasks. In our tasks, observers were cued either by spatial location or feature to two of four dot patches. They then estimated the color or motion direction of one of the cued patches, or averaged them. In all cases we found that cueing improved performance. We decomposed the effects of the cues on behavior into model parameters that separated sensitivity enhancement from sensory selection and found that both were important to explain improved performance. We found that a model which shared parameters across forms of cueing was favored by our analysis, suggesting that observers have equal sensitivity and likelihood of making selection errors whether cued by location or feature. Our perceptual data support theories in which a shared computational mechanism is re-used by all forms of attention.

**Significance Statement:** Cues about important features or locations in visual space are similar from the perspective of visual cortex, both allow relevant sensory representations to be enhanced while irrelevant ones can be ignored. Here we studied these attentional cues in an estimation task designed to separate different computational mechanisms of attention. Despite cueing observers in three different ways, to spatial locations, colors, or motion directions, we found that all cues led to similar perceptual improvements. Our results provide behavioral evidence supporting the idea that all forms of attention can be reconciled as a single repeated computational motif, re-implemented by the brain in different neural architectures for many different visual features.

## Introduction

The visual world presents human observers with an overload of sensory information, only part of which is relevant to behavioral goals at any given moment in time. Observers manage this complexity by prioritizing specific aspects of vision, such as basic features or spatial locations. These forms of attention can be operationalized by providing observers with cues about the relevance of location (Cohen & Maunsell, 2011), color (Jehee, Brady, & Tong, 2011), direction of motion (Huk & Heeger, 2000; Saenz, Buracas, & Boynton, 2002; Serences & Boynton, 2007), orientation (Rossi & Paradiso, 1995; Cohen & Maunsell, 2011) or object category (Harel, Kravitz, & Baker, 2014) among others. These cues improve response times (Eriksen & Hoffman, 1972; Posner, Snyder, & Davidson, 1980) as well as the ability of observers to detect and discriminate visual stimuli (Carrasco, 2011). Spatial and feature-based attention rely on different kinds of cues and must have their effects on different populations of neurons tuned to these visual properties. While the effects of spatial cues are local (Alvarez & Cavanagh, 2005; Cohen & Maunsell, 2011) the effects of featural ones spread across the entire visual field (Saenz et al., 2002; Saenz, Buraĉas, & Boynton, 2003; Serences & Boynton, 2007; Treue & Martinez-Trujillo, 1999; Jehee et al., 2011; Störmer, Cohen, & Alvarez, 2019; Liu & Mance, 2011; Cohen & Maunsell, 2011). The effects of both forms of cueing also appear to combine in an additive manner (Hayden & Gallant, 2009; Treue & Martinez-Trujillo, 1999; White, Rolfs, & Carrasco, 2015; Andersen, Fuchs, & Müller, 2011) with small differences in timing (Liu, Stevens, & Carrasco, 2007; Hayden & Gallant, 2005), further evidence that they operate in parallel.

Despite having their effects on different neural populations, spatial and feature-based attention may share a computational mechanism, thus providing a unified view of how different cue types enhance behavioral performance. Computational mechanisms of attention can be split into two general categories by whether they act to enhance sensory representations (Treue & Martinez-Trujillo, 1999; Luck, Chelazzi, Hillyard, & Desimone, 1997; Mitchell, Sundberg, & Reynolds, 2007; Itthipuripat, Cha, Byers, & Serences, 2017; Noudoost, Chang, Steinmetz, & Moore, 2010; Eckstein, Peterson, Pham, & Droll, 2009; Müller et al., 2006) or select and reinforce the transmission of attended features through the visual system (Desimone & Duncan, 1995; Briggs, Mangun, & Usrey, 2013; Fries, Reynolds, Rorie, & Desimone, 2001; Pestilli, Carrasco, Heeger, & Gardner, 2011; Birman & Gardner, 2019; Lee, Itti, Koch, & Braun, 1999; Pelli, 1985; Palmer, Verghese, & Pavel, 2000; Hara & Gardner, 2014). Whether spatial or featural cues use either of these mechanisms has been studied in a variety of psychophysical tasks. For example, using a masking paradigm Baldassi and Verghese (2005) showed that the effects of spatial and featural cueing are selective for properties of the mask that are congruent with the cue and suggest a model with shared mechanisms of sensitivity enhancement. Paltoglou and Neri (2012) used a psychophysical white noise paradigm and showed a similar set of results, where gain changes explained the effects of both cue types. However, Ling, Liu, and Carrasco (2009) showed using an external noise paradigm that the effects of spatial and feature-based attention do differ, but only under conditions of high noise. This finding suggests that feature and space differ in their sensitivity to irrelevant sensory information. These various findings might be reconciled by recognizing that the effects of attention differ under conditions of low and high competition. White et al. (2015) showed that the behavioral effects of spatial and feature-based cues are additive under conditions of low competition but subject to a non-linear selection effect at high competition, consistent with a two-step process of independent sensitivity enhancement followed by a selection mechanism. However, this has not been directly tested behaviorally because existing paradigms (using masking, reverse-correlation, or external noise) did not explicitly model sensitivity and selection.

Here we used a cued estimation task and modeling to test whether spatial and feature-based cues share an implementation through a combination of sensitivity enhancement and sensory selection. To do this, the form of cueing must be manipulated without a change in task or stimulus. We therefore developed a set of tasks in which cueing by location, color, and motion could be directly compared on shared metrics. We found that observers were able to use all forms of cues with similar efficacy. When broken down into sensitivity enhancement and sensory selection we found that both mechanisms played a role for spatial and feature-based cues and that a model with shared parameters for all cue types was preferred over one with separate parameters for each cue. Our results suggest that spatial and feature-based attention are more similar than different and support theories of a shared top-down computational mechanism (Treue & Martinez-Trujillo, 1999; Ni & Maunsell, 2019; Maunsell & Treue, 2006).

## Results

To measure the effect of cueing by feature or location on perceptual sensitivity we asked observers to perform a cued motion direction averaging task (Fig. 1). Briefly, observers were asked to report the average motion direction of two out of four random dot patches. The two selected patches were cued either by their common location or color, randomly interleaved across trials. This task thus engages either spatial or feature-based attention according to the current cue. In addition, the task does not require working memory, which avoids potential confounds introduced by storing perceptual information during a delay.

**Figure 1:**
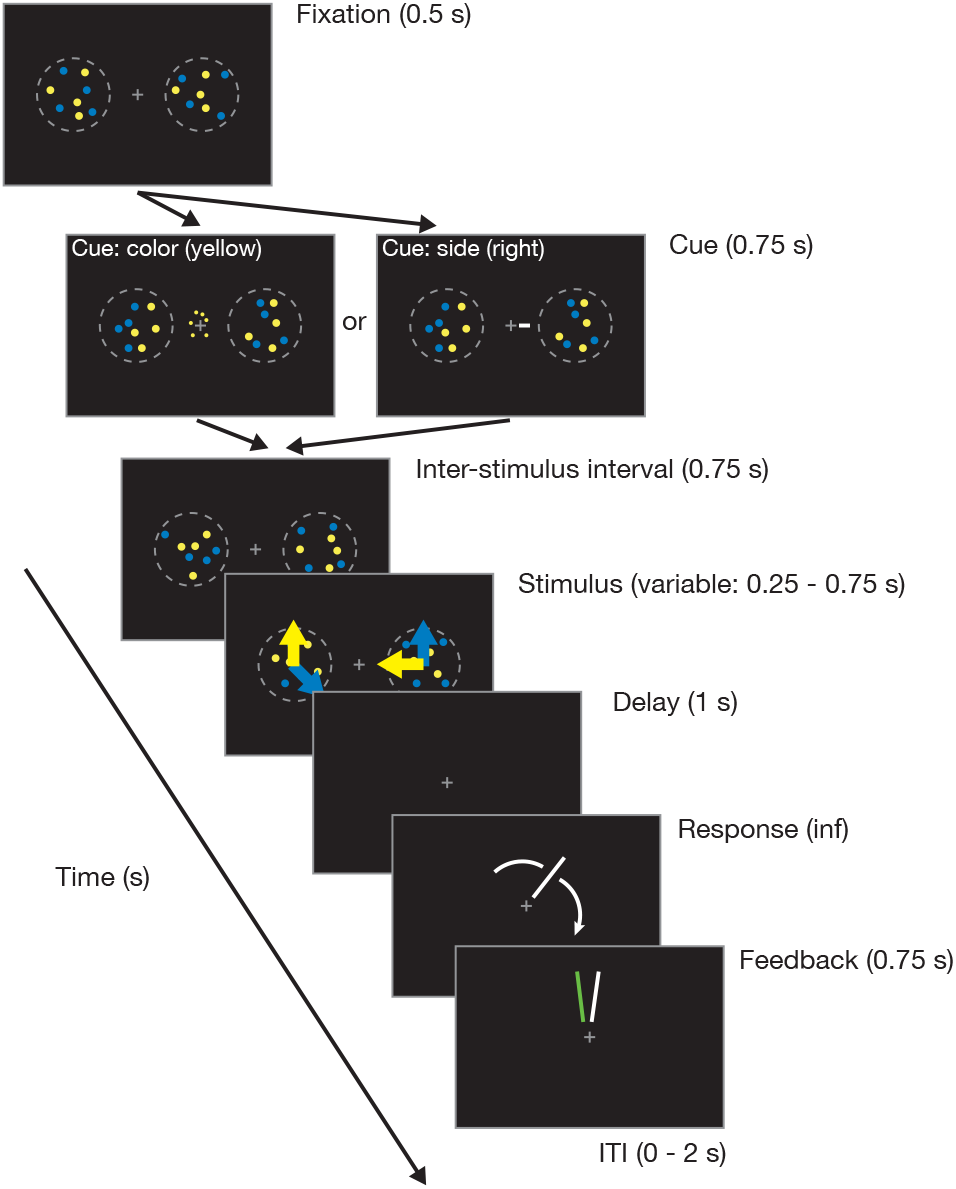
Motion direction averaging task. Observers were asked to select two out of four random dot patches and average their directions of motion. Observers initiated trials by fixating a central cross (Fixation). During this initial period and until stimulus presentation the dots in the four patches moved incoherently. A cue was shown at the fixation cross indicating which two dot patches should be averaged (Cue): a line to the left or right of fixation indicated selection by side or a mini-patch of dots colored yellow or blue indicated selection by feature. After a brief delay (Inter-stimulus interval) the four dot patches moved coherently in random directions for a variable duration (Stimulus). After another brief delay (Delay) observers used a rotating wheel to report the *average* direction of motion for the two dot patches they were asked to select. Feedback was given by indicating the true average motion direction.

By examining known stimulus manipulations we first showed that our task provided a good measure of perceptual sensitivity. As expected, we found that estimates of average direction were more precise on easier trials. This was true both for trials with a smaller angle difference between the two cued patches (Fig. 2a) and for trials with longer duration (Fig. 2b). To quantify this, we fit a model of perceptual sensitivity in estimation tasks (the “target confusability competition” model, see Schurgin, Wixted, and Brady (2020) and Methods) which fits a parameter *d′* in a manner analogous to signal detection. Increasing the stimulus duration from a mean of 0.35 s to 0.625 s made the task easier and increased the average *d′* across observers from 1.53 to 1.77 (+0.24 95% CI [0.17, 0.30]). Reducing the angle between the two dot patches from a mean of 100° to 30° also made the task easier, increasing the average *d′* from 1.48 to 1.84 (+0.36 95% CI [0.28, 0.43]).

**Figure 2:**
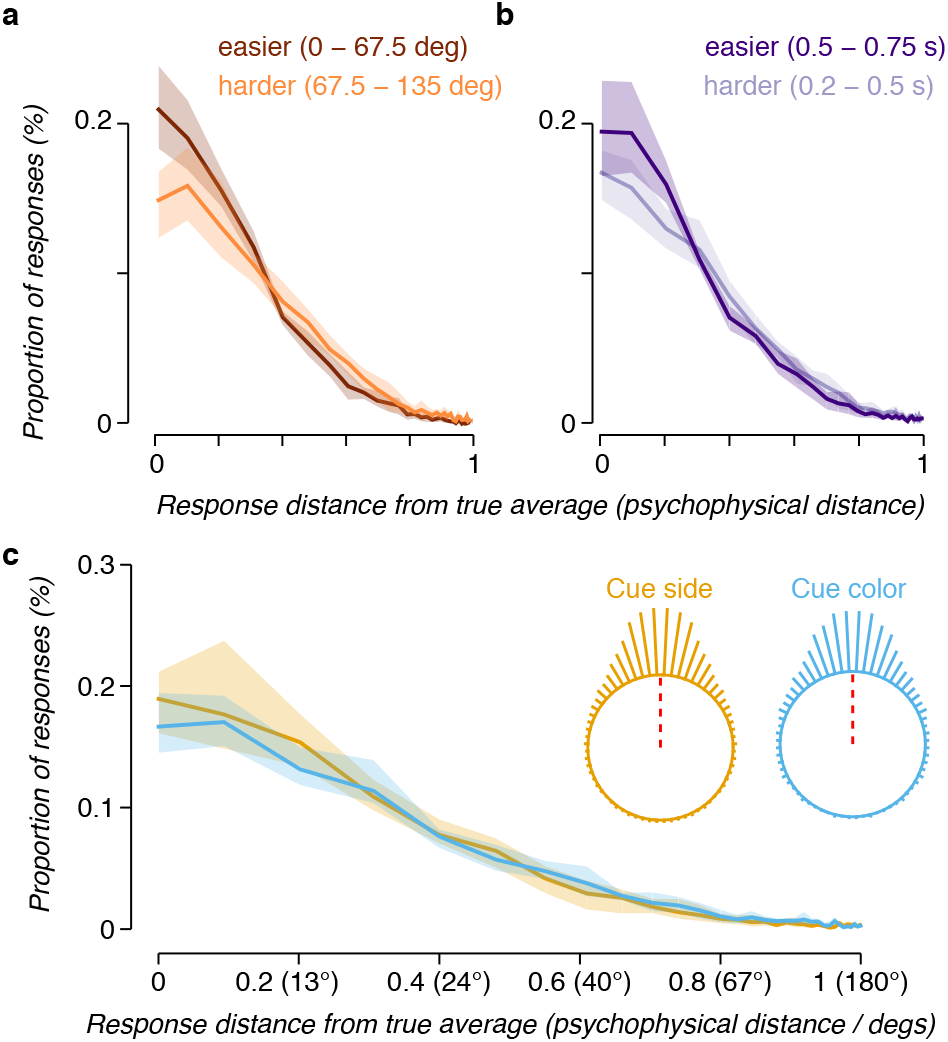
Estimation error during the averaging task. (a) A histogram displaying the proportion of responses at each absolute distance from the true average motion direction (rotated to 0) is shown, averaged across observers. Data are split by the angular distance between the motion direction of the two dot patches which were cued. Note that the x-axis in all panels has been re-scaled from degrees to psychophysical distance, see Methods. (b) Conventions as in (a), data are split by the duration of the stimulus. (c) Conventions as in previous panels. Selection by spatial location (i.e. averaging the two patches on the right or left) is shown in yellow, and selection by color (i.e. averaging the two yellow or blue patches) is shown in blue. The two inset plots show the same histogram in a circular space, with a red dashed line indicating the true average. In all panels lines indicate the average normalized histogram of response counts across observers and shaded regions the 95% confidence interval.

Having validated our perceptual sensitivity measure, we next tested whether performance was better for spatial or feature-based cues and found that spatial cues provided only a modest benefit to performance (Fig. 2c). Estimates of average direction were more accurate when observers selected dot patches by their common spatial location (on the left or right) compared to by feature (yellow or blue). The *d′* across observers was 1.72, 95% CI [1.51, 2.05] for selection by side and 1.52 [1.33, 1.69] for selection by color. This modest increase in *d′* was found for 6/7 observers, averaging 0.19 [0.08, 0.43]. Splitting *d′* out by all four selection conditions: select yellow, select blue, select left, and select right, *d′* = 1.60 [1.41, 1.73], 1.48 [1.30, 1.69], 1.73 [1.48, 1.98], and 1.75 [1.52, 2.20], respectively. Our data therefore show that changing the form of cueing has an almost negligible effect on averaging. The small advantage of spatial cueing that we did observe may in fact be explained by the known effect of stimulus distance on the accuracy of averaging (see e.g. Maule and Franklin (2015) among others).

Although the performance difference between spatial and color cues was small, the mechanism by which subjects used the cue could have been quite different. For example, cues could have improved performance by making perception more precise, or performance could also have improved if observers became more accurate in selecting the cued dot patches. These two different ways of improving performance, enhancing sensitivity and reducing selection errors, both could have resulted in similar response distributions. We next set out to separate these two computational mechanisms.

To separate changes in sensitivity from selection we devised a cued estimation task using the same stimulus as the cued averaging task (Fig. 3). In the estimation task, observers reported the properties of only a single dot patch. The advantage of this change is that a computational model can be used to separate the effects of a change in sensitivity from a change in selection. If an observer’s response is close to the angle of the target dot patch, then it is likely that the observer correctly selected the target and reported it back. If their response is instead close to the angle of one of the other dot patches, there is some likelihood that the observer made a mistake and mis-selected a non-target patch, i.e. an error in selection. This task allowed us to decompose these different possible explanations into model parameters fit to large numbers of trials.

**Figure 3:**
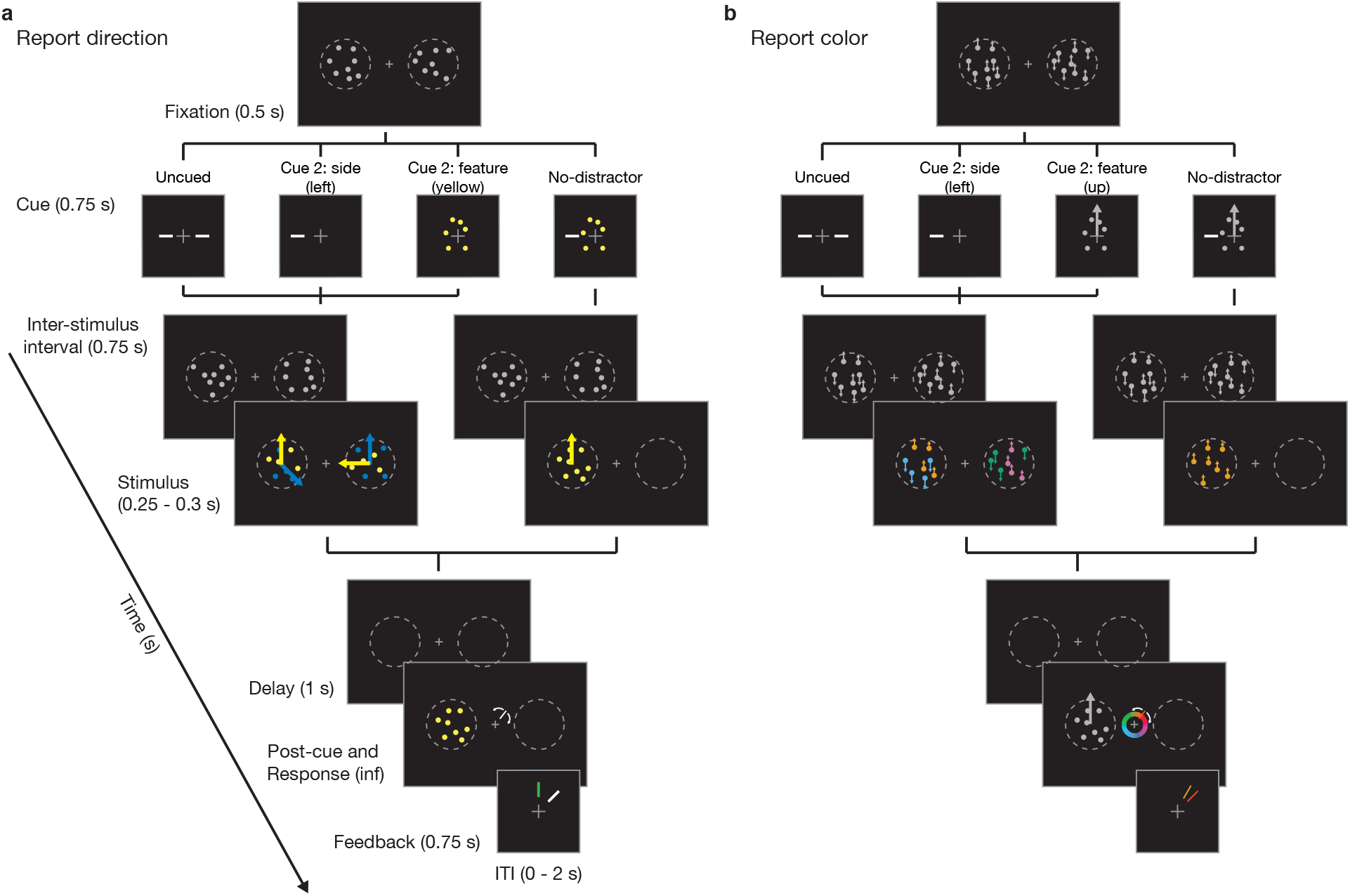
Estimation task. (a) Observers began each trial by fixating a central cross (Fixation). A pre-cue (Cue) was then shown at fixation to indicate to observers which of the four dot patches they should select. A brief delay (Inter-stimulus interval) followed. Up to this point all four dot patches were colored white and moving incoherently. The dots then became colored and coherent for a variable duration (Stimulus). After another brief delay (Delay), observers were shown a second cue which was used to disambiguate the target stimulus (Post-cue). For example, if the observer was cued to remember the two stimuli on the left, the post-cue could be yellow to indicate that of the two patches that were cued (blue and yellow, left side) only the motion direction of the target (the yellow patch on the left) should be reported. Observers were given unlimited time to respond (Response) and received feedback before the next trial (Feedback). (b) A second variant of the same task was also run in which the cues were side (left or right) or motion direction (up or down) and observers reported about the color of the dot patches.

We tested our hypothesis on two variations of the cued estimation task to ensure our results were robust to the particular cued features. In one variant observers were cued by location or color and had to report the motion direction of a dot patch (Fig. 3a). In the other, observers were cued by location or motion direction while reporting color (Fig. 3b).

We first evaluated the cued estimation task on two reference conditions: trials in which no cue was given and trials where no distractors were shown. These two references provide a lower bound (Uncued condition, Fig. 3) and an upper bound (No-distractor, Fig. 3) on performance. The no-distractor condition is an upper bound because the stimulus to report appears in the absence of distracting information. This should be equivalent to the optimal performance of an observer cued to select a single dot patch.

The reference conditions showed that observers could perform this task and that indeed, the absence of distractors improved estimates of motion direction. In both variations the observers’ estimates were less precise in the uncued condition (black markers have a wider distribution compared to grey markers, Fig. 5), indicating poorer performance. We next decomposed these responses into the separate sensitivity and selection parameters to understand what caused the deteriorated performance in the presence of distractors.

To decompose the responses in the cued estimation task we extended an existing observer model (Schurgin et al., 2020) to fit separate parameters to capture sensitivity (*d′*) and selection (*β*) (Fig. 4). On each trial, the observer model encodes the four stimulus patches (Fig. 4a,b) by a set of noisy channels (Fig. 4c). A parameter, *d′*, controls the maximum value of the channels and the sensitivity of the observer (Fig. 4d). In this model, the channel with the maximum response “wins” and becomes the observer’s estimated angle for that dot patch (red dashed line, Fig. 4e). The reported angle is then sampled from the estimated angles for the four dot patches in proportion to the *β* parameters (Fig. 4f,g). By computing the probability of each channel winning we can generate the full response likelihood for each dot patch (color distributions, Fig. 4e), i.e. how likely the observer was to make a particular response having seen that patch. When weighted by the *β* parameters, which fit the probability of choosing to report each patch, we get the distribution of response likelihoods for that trial, given all stimuli (Fig. 4h). Thus, in our model the *β* parameters control how selection occurs while the *d′* parameter controls the observer’s precision of report.

**Figure 4:**
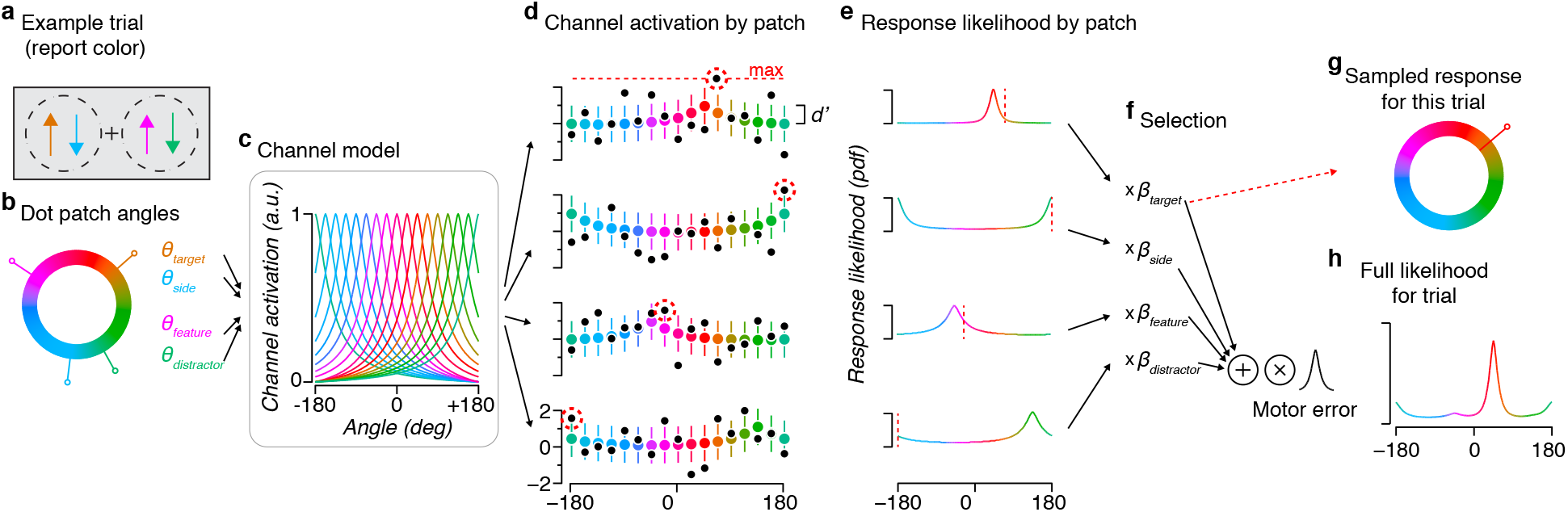
Estimation task model. (a-c) On a trial, stimuli of varying angles are encoded by many independent channels (each channel is represented by a single colored tuning curve). Each channel’s tuning profile is defined by the psychophysical distance function (see Methods) relative to that channel’s preferred angle. (d) The channel responses for a trial are noisy, so for a particular presented stimulus each channel will have a response sampled (black markers) from a normal distribution with a mean set by the height of the tuning curve at the stimulus angle (colored markers indicate mean and error bars ±1 standard deviation). The model’s predicted response to a dot patch is found by taking the channel with the maximum sampled response and reporting its preferred angle. These winning angles are shown as a red vertical dashed line in (e) along with the probability distribution of each channel having the maximum response. A free parameter *d′* sets the spread of this distribution by multiplicative scaling of the peak responses of the channels, in a manner analogous to signal detection (see Methods). (f) The selection parameters (*β*) control the probability that each dot patch will influence an observer’s report. (g) From a discrete sampling perspective, the *β* values sets the proportion of trials where the observer will report about the estimated angles of particular dot patches, e.g. here the estimated target angle is reported. (h) To fit the model we computed the full likelihood distribution for each trial. We then optimized the model parameters to maximize the likelihood of each observer’s actual reports across all trials.

We first confirmed that the model captured the perceptual sensitivity of the observers in the two reference conditions. The model accounted for the qualitative aspects of the data well (curves track the markers, Fig. 5a,b). For the report-color task the average 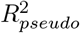 over observers was 0.84, 95% CI [0.71, 0.90] (uncued reference) and 0.88 [0.85, 0.91] (no-distractor reference). For the report-direction task 0.85 [0.74, 0.89], and 0.90 [0.88, 0.92], respectively. We also confirmed that every model we fit was a better fit to the true data than a data set in which we permuted the response array. Averaged over the two task variations the improvement in cross-validated log-likelihood for real data compared to the permuted data was 39.01, [26.43, 56.74] (uncued) and 98.60, [84.31, 115.08] (no-distractor).

**Figure 5:**
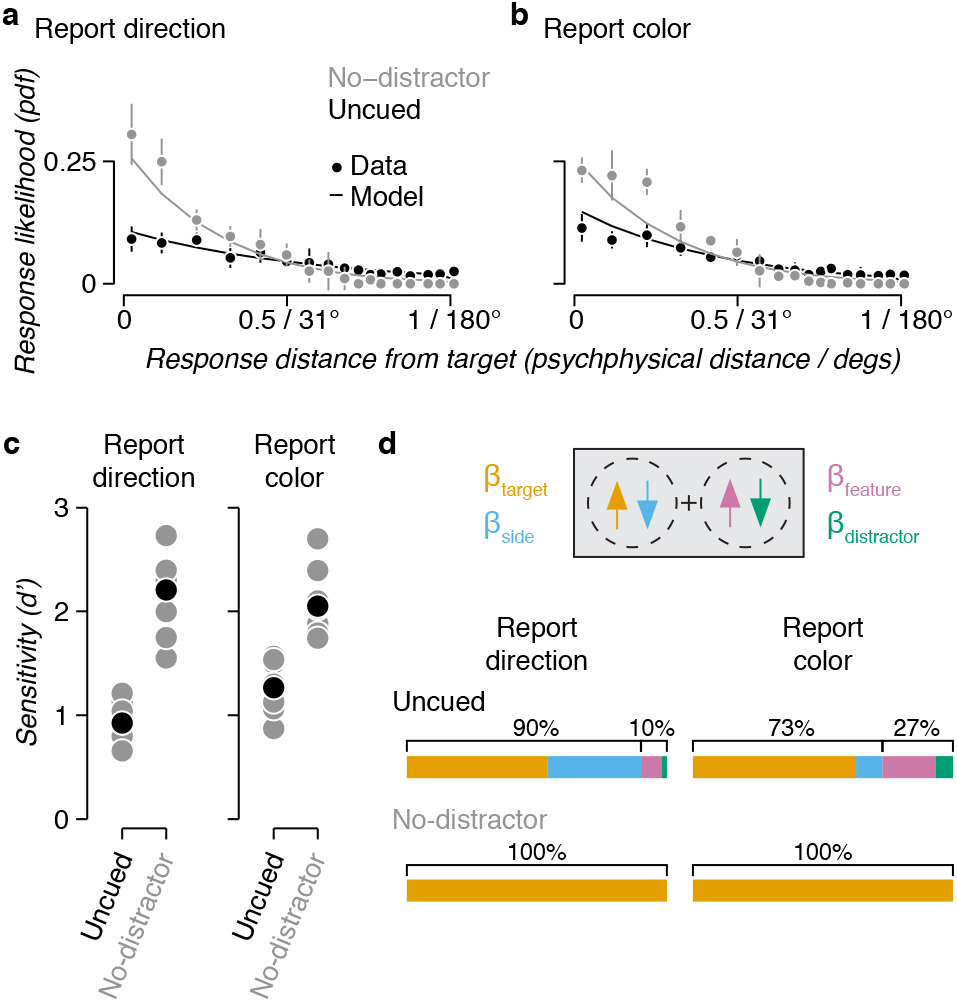
Baseline estimation performance. (a) A histogram of observer responses, relative to the true target motion direction is shown averaged across observers for the uncued and no-distractor conditions. Markers indicate the mean and error bars the 95% confidence intervals. Lines are the average fit of the model. (b) As in (a) for the report-color variant. (c) The *d′* parameter is shown for individual observers (gray) and the average (black) for each condition and task variant. (d) The *β* parameters are shown averaged across observers for each condition and task variant. *β_target_* refers to the dot patch which was post-cued (here left side, moving up), *β_side_* is the patch on the same side as the post-cued patch (left side, moving down), *β_feature_* is the patch on the other side with the matched feature (right side, moving up), and *β_distractor_* is the patch on the other side with a mis-matched feature (right side, moving down).

Looking at the sensitivity parameter we found that without distractors observers consistently made more precise reports (Fig. 5c). The average *d′* across observers for no-distractor trials in the report-direction task 2.21 [1.91, 2.46] and the report-color task was 2.05 [1.88, 2.35]. By comparison, in the presence of distractors *d′* was only 0.79 [0.58, 1.02] for the report-direction task and 1.28 [0.95, 1.49] for report-color. These differences show that adding distractors to the scene caused observers to recall the color or motion direction of a single dot patch with far more estimation error.

We also found that uncued responses were characterized by a large proportion of selection errors (all Uncued *β* values non-zero, Fig. 5d). This indicates that the presence of distractors not only reduced the accuracy of reports but caused observers to report about the incorrect patches. In the report-direction task 44% of reports were about the wrong dot patch, most often the dot patch on the same side but of the wrong color (33% [28, 38]), but also often about the feature-matched patch on the wrong side (8% [2, 17]). In the report-color task observers made 35% of reports about the wrong dot patch. Most often about the patch on the same side (19% [12, 31]) but sometimes about the feature matched patch (6% [1, 12]) or the distractor (10% [4, 19]). In summary, adding distractors decreased the precision of estimates and caused observers to report about the entirely wrong dot patch for about one in every three trials.

Returning to the main hypothesis, we next looked at whether the cued trials caused a change in sensitivity or selection when compared to baseline performance. In the cued trials, observers selected dot patches by their common spatial location (left or right) or common feature (yellow/blue color, or up/down motion for the two task variants, respectively) (Fig. 3). Although cued to two dot patches, observers still reported only the properties of a single dot patch at the end of each trial, uniquely identified by the post-cue. As expected, we found performance on these trials to be intermediate between the reference conditions (Fig. 6a,b). Again, we confirmed that the model fits explained a substantial portion of the variance, finding an 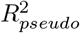 of 0.90 [0.86, 0.93], 0.89 [0.88, 0.91] for the report-direction variant and 0.91 [0.88, 0.92] (spatial cued), 0.87 [0.80, 0.92] for the report-color variant. We also confirmed that the models far exceeded the fit to permuted data sets, with an average increase in cross-validated likelihood of 231.49, [162.04, 315.85].

**Figure 6:**
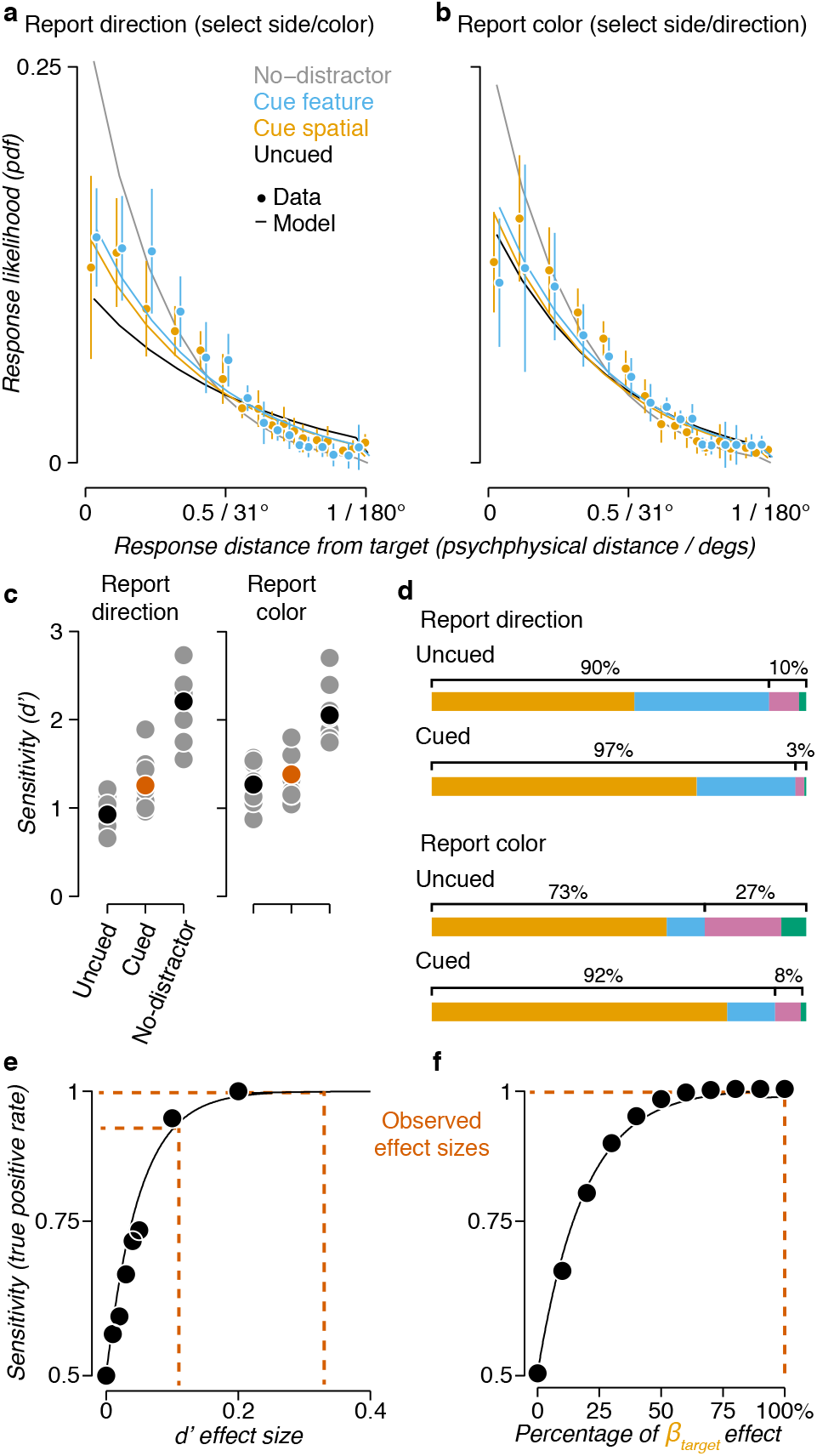
Comparison of spatial and feature-based cueing. (a) A histogram of observer responses, relative to the true target motion direction, is shown averaged across observers. Markers indicate the mean and error bars the 95% confidence interval. Estimates are split by whether the pre-cue indicated a feature (Cue feature, blue) or a spatial location (Cue spatial, yellow). (b) As in (a) for the report color variant. (c) The value of the *d′* parameter is shown for the cued data collapsed across the two cues, compared to the uncued and no-distractor conditions. (d) The value of the *β* parameters are shown for the cued data, again collapsed across the two cues. Refer to Figure 5d for legend. (e) Results of a power analysis simulation for the *d′* parameter. Markers indicate actual simulated datasets and the line is the fit of an exponential function. Red dashed lines indicate the effect sizes reported in the paper. (f) As in (e), for the results of a power analysis of the *β* parameter. Note that the x-axis represents the percent difference between the 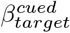 and 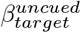, so the reported effect size is 100%.

Although both the uncued and cued trials had distractors present, we found that cueing reduced the impact of the distractors, both improving sensitivity and reducing selection errors. To quantify this, we combined trials from the two cued conditions and looked at the values of *d′* (Fig. 6c) and *β* (Fig. 6d). Cueing, by spatial location or feature, increased *d′* in both task variants (Fig. 6c). This increase was modest, the uncued *d′* was 0.93 [0.79, 1.05] and 1.27 [1.10, 1.44] for the report-direction and report-color variants, respectively. In the cued model these were 1.26 [1.09, 1.51] and 1.38 [1.21, 1.62]. Taking the uncued condition as a baseline and the no-distractor condition as ceiling, this improved sensitivity corresponds to 23.4% and 17.5% of the no-distractor increase. We found that a model with separate *d′* parameters for cued and uncued trials better accounted for our data compared to a model which combined these trial types: the average increase in cross-validated likelihood was 2.82 [0.82, 5.06] for report-direction and 6.70 [3.52, 11.73] for report-color. We also observed substantial reduction in errors of selection. Comparing uncued to cued, observers increased their selection of the target dot patch by 15% [9, 23] and reduced incorrect selection of the same-side by 7% [−14, −1], same-feature by 6% [−12, −2], and distractor by 2% [−7, 3] for the report color variant. For the report direction variant we found that observers increased target selection by 18% [8, 25], same-side decreased by 6% [−17, 2], same-feature by 3% [−9, 0] and distractor by 9% [−18, −3]. Again, we found that a model with separate *β* parameters for cued and uncued trials better accounted for our data: the average increase in cross-validated likelihood was 2.89 [0.10, 5.74] for report-direction and 4.84 [1.15, 10.20] for report-color. These data show that cueing has a substantial impact on performance in this task, improving sensitivity and reducing the probability that observers will misreport about the irrelevant dot patches.

Because the change in *d′* and *β* that we observed were small we performed a model recovery simulation to confirm that our dataset was sufficient to detect these effects with high power (Fig. 6e,f). Briefly, we simulated data sets with various *d′* and *β* parameters using the range of values observed in the cued and uncued conditions (Fig. 6c,d). We generated 200 such data sets for every combination of parameters and fit these with our analysis pipeline. We then bootstrapped the resulting values to compute the model’s true positive rate (sensitivity), i.e. the probability that the model would recover a difference in *d′* or *β* when a real difference was simulated with noise (see Methods: Model recovery for details). We found that the effects we observed were well within the range that we would expect to be able to detect (observed effect sizes are all above 90% sensitivity, Fig. 6e,f). While the averaging task data showed that human observers could use spatial and featural cues with similar efficacy, this could be done by similar or different computational mechanisms. Testing this hypothesis by fitting our model with either shared or separate parameters for the Uncued, Cue spatial, and Cue feature conditions (Fig. 7a), revealed that spatial and featural cues employ a shared computational mechanism. We first fit a Null model which used the same parameters for all three conditions, testing the null hypothesis that cueing had no effect. We then compared this to a Cued model which separated the spatial and feature cueing conditions from the uncued condition (the parameters from this model are reported in Fig. 6c,d). Finally, we fit a model in which all three conditions had separate parameters. We found that separately fitting cued trials improved model fit (Fig. 7b), but there was no additional improvement to fitting spatial and feature-based conditions separately. Adding separate cued parameters increased the cross-validated likelihood by 4.11 [1.14, 7.34] and 8.69 [4.17, 13.93] for report direction and color, respectively. Further separating spatial and feature-based trials did not further improve the fits: change in average cross-validated likelihood 3.28 [−0.49, 6.21] and 0.71 [−2.22, 7.50]. Note that real, but small, differences between spatial and feature cueing would be unlikely to have been detected at our power (i.e. a difference in *d′* < 0.05 or *β* < 0.01 Fig. 6e,f). In summary, because the improvements in perceptual sensitivity and reduction in selection errors were shared between cueing by location, color, and motion direction, our results suggest that spatial and feature-based selection share a common computational mechanism that both enhances sensitivity and reduces errors of selection.

**Figure 7:**
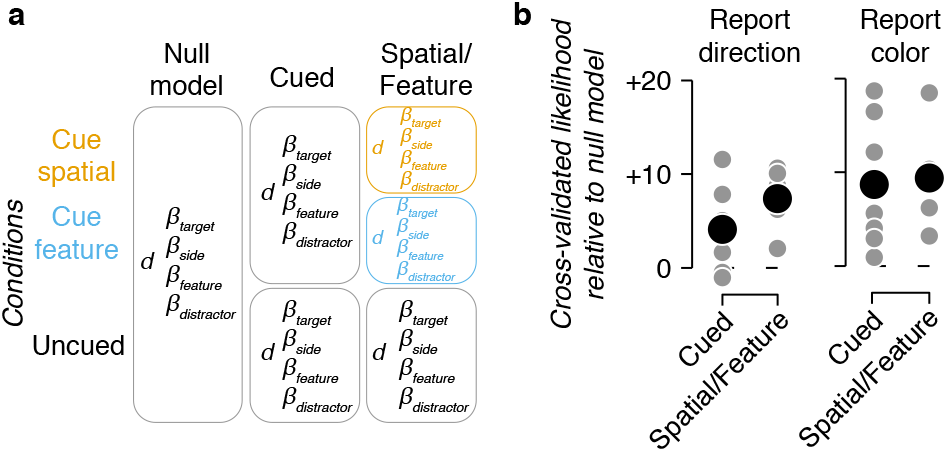
Spatial and feature-based cueing share model parameters (a) Diagram showing how parameters were fit for the three models we compared. Each row are the trials for the two cued conditions and the uncued condition. Each column shows how parameters were fit to the different sets of data. (b) The relative cross-validated likelihood of the Cued and Spatial/Feature model are shown compared to the Shared model for individual observers (grey markers) and the average (black marker). Some markers are hidden by others.

## Discussion

We found that spatial and feature-based cues changed perception in similar ways across a set of cued estimation tasks. In one task observers averaged motion directions cued either by location or color, a perceptual judgment with low working memory load. We found that observers were able to use both cues with similar efficacy, suggesting that they had similar effects on sensitivity. Although this measurement put each cue on the same scale it did not demonstrate computational similarity: observers might, for example, use sensory enhancement more when cued to a spatial location. We designed a set of tasks to separate sensitivity enhancement from errors of selection through the use of estimation (Prinzmetal, Nwachuku, Bodanski, Blumenfeld, & Shimizu, 1997; Prinzmetal, Amiri, Allen, & Edwards, 1998) and a computational model. Our data showed that all cues improved perceptual sensitivity and reduced errors of selection. Model comparison revealed that a model with shared parameters for spatial and feature-based cues was preferred over models in which perceptual improvements were separated by cue type. In other words, in our experiments the exact nature of the cue, whether spatial location, color, or motion direction, did not change the effects.

Several previous studies have also looked at how perceptual data inform us about the mechanisms of spatial and feature-based attention. Although many of these studies used different tasks or stimuli between cueing conditions we can nevertheless make indirect comparisons about the underlying computational mechanisms. For example, Ling et al. (2009) asked observers to judge motion directions under varying amounts of noise. They showed that at low noise both spatial and featural cues improved performance, but only featural cues improved performance in the presence of high external noise. The low noise comparison confirms that sensitivity enhancement is shared between different forms of cueing (Martinez-Trujillo & Treue, 2004; Cohen & Maunsell, 2011). Although the authors reported differences between cues under high noise conditions, this may be the result of using the same feature dimension for cue, report, and noise. Observers were cued to a motion direction, then reported whether a stimulus was rotated relative to the cue under conditions of decreasing motion coherence. In general, changes in tuning functions reappear whenever feature-based cues are matched with the dimension of report (Ling et al., 2009; Paltoglou & Neri, 2012; Baldassi & Verghese, 2005) while gain combined with a sensory selection step, as we propose, is sufficient to explain attentional effects cued by orthogonal features (White et al., 2015; Ni & Maunsell, 2019) such as those in our tasks.

In another comparison, Baldassi and Verghese (2005) measured perceptual thresholds for oriented stimuli in the presence of masks. These masks varied in their distance and similarity to the target, defining a set of tuning functions. They found that an exogenous location cue at the target improved perception regardless of the mask distance or difference from the target, while exogenous orientation cues only provided a benefit when the mask was close to or similar to the detected orientation. Their data and ours are consistent with models in which tuned filters increase their precision according to their similarity to the cue (Treue & Martinez-Trujillo, 1999). Their data also points to the possibility that exogenous cueing relies on a different set of mechanisms that may be specific to spatial cues (Busse, Katzner, & Treue, 2006; Donovan, Zhou, & Carrasco, 2020).

Several previous studies used task designs in which a direct comparison was possible. In one, Liu et al. (2007) looked for differences in timing between spatial and featural cueing. They found a small advantage to spatial cueing when the cue preceded the stimulus by 0.2 s, with both forms of cueing becoming equivalent at a 0.5 s delay. This result suggests that spatial attention can be deployed faster than featural attention (Goddard, Carlson, & Woolgar, 2019), a finding that explains in part why spatial attention is sometimes considered “primary” over featural attention (Wolfe, 1994). We interpret the faster deployment of spatial attention in the same way the authors do, as a difference in the neural architecture of top-down attentional control. Timing differences alone are insufficient to suggest that computational differences exist between spatial and featural attention. Because the performance enhancement of cueing is otherwise identical, both our results and theirs support the possibility of similar computational mechanisms.

Another direct comparison was performed by White et al. (2015). The authors looked for evidence of independent mechanisms by examining whether location cues and featural cues combine in an additive manner. In a low-competition condition they found that cues combined in a manner consistent with two independent additive mechanisms. In a high competition condition they found that a second selection step was necessary to explain the effects of combined cues. They perform this step in their model by divisive normalization. Our model used proportional sampling as a way to approximate these more complex biophysical mechanisms, but the end result is the same: there is a winner-take-all effect in which the stronger cued responses are more likely to out-compete weaker uncued ones.

In the cued estimation task we used a post-cue to reveal the target in each trial. This afforded us the flexibility to cue different dimensions of the stimuli while still allowing us to measure an observer’s knowledge about any of the stimuli on the screen, providing us with the data necessary to separate sensitivity from selection. To avoid observers treating cued and uncued targets differently (Rahnev et al., 2011) we chose not to include invalid targets, i.e. we never asked the observers to report about a dot patch that was not part of the pre-cued set. One consequence of this is that we don’t know how much information observers retained about stimuli in the absence of attention. Many previous reports about dual task performance (Lee et al., 1999; Reddy, Wilken, & Koch, 2004; Lee, Koch, & Braun, 1997) and unattended stimuli (Li, VanRullen, Koch, & Perona, 2002; Birman & Gardner, 2019) show that visual information is processed and retained even at low levels or in the absence of attention. What our design did allow us to measure was that errors of selection decrease with attention. This is consistent with models in which some stimuli are prioritized for processing over others (Baldassi & Verghese, 2005; Ling et al., 2009; White et al., 2015) but does not preclude the possibility that information is retained about all stimuli.

One alternative to our model is that observers might encode an “ensemble” representation of the stimulus (Utochkin & Brady, 2020) instead of encoding the four dot patches independently and sampling from them (Emrich & Ferber, 2012; Bays, Catalao, & Husain, 2009; Bays, Wu, & Husain, 2011). Some researchers have shown evidence for ensemble representations by measuring bias toward the mean of large sets of stimuli (Utochkin & Brady, 2020; Brady & Alvarez, 2011). In the task used by Utochkin and Brady (2020) stimulus angles are sampled from a distribution with an informative mean, making an ensemble representation an optimal strategy. The mean angle in our task was uninformative across trials, even though on a small number of trials with clustered stimuli it could have provided useful information. This key difference, if recognized by observers, likely played a role in which strategy they used to solve each task. In addition, our model represents the ensemble statistics in an implicit manner because of how the channels encode the stimuli. To see this, consider Figure 4 for a trial with four dot patches of similar color. The channel responses would all peak at nearby values, causing an ensemble-like effect in which the mean angle becomes more likely to be reported than any individual dot patches’ true color angle. We expect that this behavior in the model should account for the effect reported by Utochkin and Brady (2020), but an ideal comparison could be performed by designing a stimulus set in which the ensemble is either informative or uninformative across different blocks of trials.

Our results are consistent with a single shared computational mechanism of attention implemented by a different neural architecture depending on the feature (Cohen & Maunsell, 2011). This view is supported by the similarity of neural effects for different forms of attention (Cohen & Maunsell, 2011; Treue & Martinez-Trujillo, 1999; Patzwahl & Treue, 2009; Ling, Jehee, & Pestilli, 2015; Jehee et al., 2011; Martinez-Trujillo & Treue, 2004), the additive or multiplicative advantage of combining multiple cues (Hayden & Gallant, 2009; White et al., 2015; Andersen et al., 2011; Goddard et al., 2019), and the similar top-down sources in prefrontal cortex from which attention signals are thought to originate (Corbetta & Shulman, 2002; Moore & Armstrong, 2003; Zhou & Desimone, 2011; Bichot, Xu, Ghadooshahy, Williams, & Desimone, 2019; Liu & Hou, 2013). From this perspective, spatial location is just another feature and spatial attention is a special form of feature-based attention (Treue & Martinez-Trujillo, 1999). Like color or motion direction, location is a dimension in the possible space of stimulus properties and neuron tuning. Although all features are treated identically in most ways, the topological representation of space in early visual cortex leads to one important difference: any local computation, e.g. response normalization (Carandini & Heeger, 2011), will have spatially-tuned effects. Consistent with this, Ni and Maunsell (2019) recently showed that the neural effects of different cue types are consistent with a common top-down mechanism combined with spatial response normalization.

A common mechanism combined with normalization predicts that the shift in spatial attention in our task, between selecting two overlapped dot patches or two separated patches, will lead to a change in the pool of activity that drives normalization early in visual cortex. Changes in the relative size of stimuli and the normalization pool are known to affect performance for low level features such as contrast discrimination (Herrmann, Montaser-Kouhsari, Carrasco, & Heeger, 2010). Other work has shown that for overlapping stimuli, as in our design, contrast normalization in early visual cortex leads to a biased representation of lower-level features but this bias does not occur for higher-level features such as motion direction (Wiesner, Baumgart, & Huang, 2020). This is consistent with our results, where we showed no difference in performance despite a change in the size of the normalization pool (from two dot patches for spatial cues to four for featural cues) and confirms that the representations of higher-level features are not affected by spatial normalization in the same manner as low-level ones.

Researchers studying visual search have long held that visual features are extracted and processed in a parallel step where spatial information is prioritized (Treisman & Gelade, 1980; Wolfe, 1994). Physiology experiments, in turn, have gone on to separate the neural effects of spatial and feature cues using different tasks. Because of these operational differences, many studies have found that spatial and featural cues have unique behavioral and neural properties—while a parallel literature of computational models (Treue & Martinez-Trujillo, 1999; Ni & Maunsell, 2019) has shown that all forms of cueing can be reconciled. The similar behavioral effects of cueing location, direction of motion, and color in our experiments demonstrate that perceptual data also provide support for a single shared mechanism of attention.

## Methods

### Observers

In total 16 observers were subjects for the experiments (9 female, 7 male, mean age 25 y, range 19 - 37). All observers except one (who was an author) were naïve to the intent of the experiments. Three observers were excluded during the initial training sessions and one after data collection due to an inability to maintain appropriate fixation (see eye-tracking below). Potential observers were not considered for inclusion in the study if they self-reported any anomaly of color vision (e.g. color-blindness). At the start of the experiment observers completed the Ishihara test for color vision (Ishihara, 1987) and one observer was excluded due to anomalous responses. Observers wore lenses to correct vision to normal if needed. Procedures were approved in advance by the Stanford Institutional Review Board on human participants research and all observers gave prior written informed consent before participating.

Seven of the observers completed the estimation task, performing on average 1428 trials (range 880 - 2123) in two to four 60 minute sessions. Seven of the observers performed the averaging task, completing on average 1010 trials (range 280 - 1475) in two to four 90 minute sessions. One observer participated in both tasks. Observers were trained for one hour on their first day and then performed at most two sessions on subsequent days, returning several times to complete the experiment.

### Hardware setup for stimulus and task control

Visual stimuli were generated using MATLAB (The Mathworks, Inc.) and MGL (Gardner, Merriam, Schluppeck, & Larsson, 2018). Stimuli were displayed on a 22.5 inch VIEWPixx LCD display (resolution of 1900×1200, refresh-rate of 120 Hz) at a 60 cm viewing distance. Output luminance and spectral luminance distributions were measured for the LCD display with a PR650 spectrometer (Photo Research, Inc.). The gamma table for the display was adjusted to linearize the output luminance separately for each color channel. The luminance spectra of the monitor was used to compute a transformation matrix from the CIELAB color space to the RGB output of the screen, such that the a* and b* dimensions could be separately manipulated without changing the luminance (L*) (C.I.E., 1978). Experiments were performed in a darkened room where extraneous sources of light were minimized. Observers used a rotating response device to provide their responses (Powermate USB, Griffin Audio).

### Eye tracking

Eye-tracking was performed using an infrared video-based eye-tracker at 500 Hz (Eyelink 1000; SR Research). Calibration was performed at the start of each session to get a validation accuracy of less than 1 degree average offset from expected, using a thirteen-point calibration procedure. Calibrations were repeated as needed after breaks. During training, trials were initiated by fixating the central cross for 0.5 s and canceled on-line when an observer’s eye position moved more than 1.5 degree away from the center of the fixation cross for more than 0.3s. Observers were excluded prior to data collection if we were unable to calibrate the eye tracker to an error of less than 1 degree of visual angle or if their canceled trial rate did not drop to near zero. During data collection the online cancellation was disabled and trials were excluded if observers made saccades outside of fixation (> 1.5deg) during the stimulus period.

### Experimental design

Stimuli for both the averaging and estimation task consisted of two pairs of overlapped dot patches, to the left and right of a central fixation cross (0.5 × 0.5 deg). The dot patches were circular regions centered 8 degrees eccentric with a diameter of 10 deg, covering from ±3 to ±13 deg along the horizontal axis and ±5 deg along the vertical axis. Each circular region contained two dot patches which differed in color and motion direction. Dots within a patch (0.3 deg diameter, 0.2 dots / deg^2^) were given an identical color and moved in the same direction at 3.5 deg / s. Each dot had a lifetime of 0.25 s, before being redrawn at a new random location. In the averaging task one patch on each side was colored yellow and one blue (90 deg and 270 deg, in a* b* space, with L* = 60).

#### Averaging task

On each trial in the averaging task observers were asked to report the average motion direction of two dot patches (Fig. 1). Before stimulus presentation a cue indicated to observers the features they would use to select the two dot patches. There were two ways that observers were instructed to select these dot patches out of the four on the screen: they could either be the two on the same side (left/right) or the two with the same color (yellow/blue). Observers were instructed at the beginning of each block of 20 trials about which form of selection would be cued, by the phrase “cue side” or “cue color”. In cue-side blocks, a line to the left or right directed observers to average the two dot patches on the corresponding side. In cue-color blocks, a miniature patch (0.2 dots / deg^2^, 0.1 deg diameter) of yellow or blue colored dots directed them to average those patches. The cues thus uniquely identified the pair of dot patches that the observer needed to select and report. Each trial was initiated by the observer fixating the central cross for 0.5 s. This was followed by the 0.75 s cue. After a 0.75 s delay the stimulus was shown.

During the stimulus the four dot patches moved coherently in random directions. The two cued patches were constrained to move in directions that were less than 135 degrees apart. This avoids response confusion because for two patches that are 180 degrees apart there are two possible responses. Observers were shown the stimulus for 0.25 - 0.75 s (randomly sampled from a uniform distribution), then allowed unlimited time to rotate the response wheel and click it to make a response. Response direction was indicated by a small line, which rotated as the observer turned the wheel. Feedback was given by showing the actual average motion direction (small green line, Fig. 1). Each trial was followed by a brief inter-trial interval (0 - 2 s, uniformly distributed).

#### Estimation task

On each trial in the estimation task observers were asked to report about either the color or motion direction of a single dot patch (Fig. 3). Before each block of 40 trials, observers were told which feature would be reported with either the phrase “report color” or “report direction” appearing on the screen. On each side during report-direction blocks one dot patch was colored blue and the other yellow and all four dot patches moved in random directions (0 to 359 deg, uniformly distributed). During report-color blocks the stimulus properties were inverted. On each side one dot patch moved upwards and the other downwards and the four dot patches were colored using angles in L*a*b* space (L* = 60, a*= cos *θ*, b*= sin *θ*, *θ* sampled from 0 to 359 deg, uniformly distributed).

Before stimulus presentation a pre-cue indicated to observers the features of the target patch or gave no information, depending on the type of trial. There were two ways that observers were instructed to select dot patches out of the four on the screen: they could either be the two on the same side (left or right) or the two with the same feature (yellow or blue in cue-color blocks, up or down in cue-direction blocks).

These pre-cues were the same as in the averaging task, either lines (cueing left or right) or patches of dots (cueing either color or motion direction). The pre-cues were blocked, so that the same cue type was repeated for twenty trials (i.e. cue side trials repeated, sampled randomly between cue left and cue right). A post-cue always indicated the specific patch that needed to be reported. Each trial consisted of the following sequence (Fig. 3): a fixation period (0.5 s), a pre-cue (0.75 s), an inter-stimulus interval (0.75 s), stimulus presentation (0.25 or 0.3 s), a delay (1 s), a post-cue resolving which dot patch should be reported (0.75 s) and then unlimited time to report a response. The inter-trial interval was 0 - 2 s, uniformly distributed. The stimulus duration (0.25-0.3 s) was chosen based on the averaging task to make the estimation task difficult for participants.

We also included uncued and no-distractor conditions as references. Comparing cued to uncued trials let us test for improved performance due to cueing, while trials without distractors gave us a measurement of the performance ceiling. In the uncued condition (Uncued, Fig. 3) observers were shown an uninformative pre-cue, then shown a post-cue which resolved which dot patch should be reported. In the no-distractor condition only the target patch was shown and observers reproduced the color or motion direction without interference. These control conditions were otherwise identical in timing to the regular trials and were also blocked in twenty trial sets. In total, 30% of trials were cue side, 30% cue feature, 20% uncued, 10% no-distractor, and an unused condition accounted for the last 10%.

### Statistical Analyses

#### Psychophysical distance

We designed our analyses to avoid conflating poor sensitivity with high lapse rates by converting angular to psychophysical distance (Schurgin et al., 2020). When observers estimate motion direction or color in angular space, including in our data, they often make a large number of responses far from the target angle. At first glance, these appear to be guesses on lapse trials (Zhang & Luck, 2008). Previous work has demonstrated that instead observers are making these low-probability responses with high confidence (Schurgin et al., 2020; Bays, 2014). This result is consistent with a continuous view of working memory (Ma, Husain, & Bays, 2014; Taylor & Bays, 2020) and can be explained by recognizing that observers are not encoding angles in degree space but in an unknown internal representation. In this representation, which we refer to as psychophysical space, angular distances that are far apart or very similar are compressed. We approximate this with a sigmoidal function:

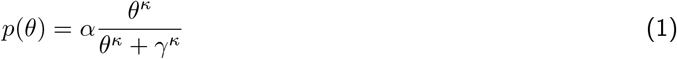

This equation transforms an angular distance *θ* to the normalized psychophysical distance *p*(*θ*), measured in perceptual units. The free parameters controlling the shape (*α* = 1.1, *κ* = 1.5, and *γ* = 35) were set according to data available in Schurgin et al. (2020) and the results are robust to small changes (on the order of 20%) in the parameters.

The parameters set above imply that an observer perceives the difference between *p* = 0 and *p* = 0.5 (*x* = 0 and *x* = 31 deg, respectively) as equal to the difference between *p* = 0.5 and *p* = 1 (*x* = 31 and *x* = 180 deg). In this way, the psychophysical space is approximately a log compression of the original degree space.

#### Averaging task analysis

To quantify the observer accuracy in the averaging task we fit a simple model of perceptual sensitivity for angular estimation tasks, the “target confusability competition” model (Schurgin et al., 2020). In the following sections we will build up this model of observer behavior.

The model takes into account two aspects of sensory representations to predict observer behavior. First, the model takes into account the “confusability” of stimuli by transforming angular distances into psychophysical distance (Eq. 1). In a second step, noisy internal channels tuned according to the psychophysical distance independently “compete” to represent a stimulus in a manner analogous to signal detection (Fig. 4a). On each trial the model proceeds according to the following steps.

First, the stimulus angles are encoded by the channels. The tuning profile of each channel takes the form of the normalized psychophysical distance function (Eq. 1). An example from the estimation task is shown in Figure 4. For a single trial with four dot patches (Fig. 4a) of varying color (*θ_target_*, *θ_side_*, *θ_feature_*, and *θ_distractor_*, Fig. 4b) a small set of channels (Fig. 4c) would be activated as in Figure 4d. The mean activation and range of noise are shown (colored markers and error bars) as well as examples of samples from those distributions (black markers). We use 100 channels in the full model, but the exact number of channels is an arbitrary hyperparameter. More channels provide better resolution to account for the data, up to some ceiling. In simulations we found that more than 100 provided a marginal benefit because correlations quickly accumulate in nearby channels.

Each channel’s response (Fig. 4d) is normally distributed around the mean (*μ*) determined from the tuning profile with standard deviation (*σ*) set to one:

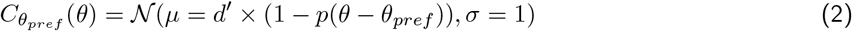

Where *θ_pref_* is the preferred orientation for that channel, *p* is the function described in Eq. 1, and *d′* controls the maximum amplitude of the response.

Next, we take the channel with the maximum response. The preferred orientation of this channel is the angle reported by the modeled observer (Fig. 4d). Because each channel has independent normally-distributed noise, the probability of any channel being reported can be computed as the conditional probability of that channel exceeding all of the other channels (Fig. 4e). We approximate this distribution by numerically integrating over channel responses *a*:

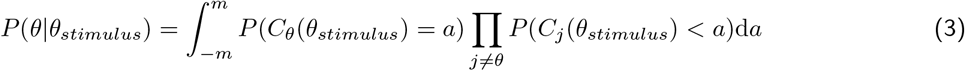

This equation computes the probability that channel *C_θ_*’s response will exceed all the other channels and be chosen as the observer’s response, given that they observed a dot patch with angle *θ_stimulus_*. *a* indexes the response of the channels according to Eq. 2. To compute the likelihood distribution across all angles we numerically evaluate Eq. 3 for each channel. We evaluate *a* in the range based 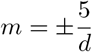 on simulations which showed that this range was more than sufficient to capture the range of channel response values, but still be computationally tractable. The likelihood distributions are normalized as probability density functions, such that:

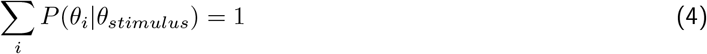

A free parameter *d′* controls the maximum amplitude of the channel responses (Fig. 4d) and therefore the width of the likelihood distributions.

This model behaves in a manner analogous to signal detection. In the simplified case of a 2-AFC task the entire model simplifies to signal detection. In such a task, an observer might be looking for a dot patch moving at an angle *θ* = 0. On each trial two items would be presented: for example, one dot patch moving in the target direction *θ* = 0 and a second in the opposite direction *θ* = 180, or *p*(*θ*) = 0 and *p*(*θ*) = 1, respectively. The channel corresponding to the target direction, *C*0 would then have a response sampled from a normal distribution according to the response to each dot patch. If *d′* = 1 these would be 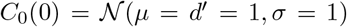 and 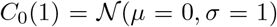. In this simplified scenario with only a single channel the *d′* parameter is equivalent to signal detection: 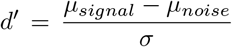, since it scales the distance between two normal distributions with *σ* = 1. The full model has 100 channels that are correlated to each other (due to the tuning functions), but the analogous behavior to signal detection holds.

For the averaging task we fit *d′* to the responses of individual observers by maximizing the likelihood of the observed data using Bayesian adaptive direct search (Acerbi & Ma, 2017). Note that the *β* parameters are used to fit the estimation task. For the averaging task we simply set *β_target_* = 1 and the others to 0. We cross-validated the models by separating the data into ten folds using nine to fit the model and evaluating the likelihood on the left out fold. We repeated this leave-one-fold-out procedure to obtain the likelihood of the full dataset.

To account for motor error we convolved the likelihood functions (Eqn. 3) with an additional 2*◦* full-width half maximum normal distribution (Fig. 4f). We also tested models with 1 and 3° distributions to ensure the results were robust to this parameter.

#### Estimation task analysis

To understand how observers encoded the stimulus during the estimation task, we expanded the model to separate sensitivity (how precise an observer’s reports were) from errors of selection (how likely observers were to report about the target or an erroneous patch). The estimation task model generalizes the averaging task model to account for the presence of four stimuli, allowing all four to modify an observer’s reports.

To model the observer’s trial-by-trial response, we assumed that four likelihood distributions (one for each of the four dot patches) were sampled according to different probabilities. The dot patches shared a sensitivity parameter (*d′*, Fig. 4d). We then modeled responses as probabilistic samples from the four distributions according to a set of bias (*β*) parameters:

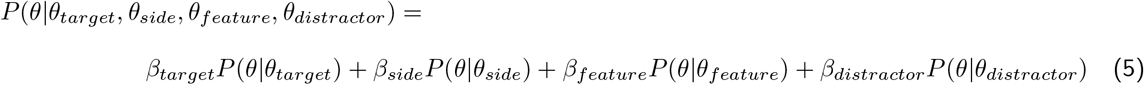

*P* refers to the probability distribution (Eq. 3) where *θ* are the possible response angles and the subscripted *θ* parameters are the angle of each dot patch (either motion direction or angle in color space). The subscript terms *target*, *side*, *feature*, and *distractor* correspond to the dot patch that was post-cued on the trial (orange), the patch on the same side (blue), the patch on the opposite-side with matched-feature (pink), and the patch on the opposite-side with mismatched-feature (green), respectively (Inset panel, top right, Fig. 4a,b).

The actual bias (*β*) values were calculated from three intermediate values:

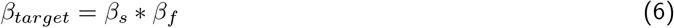

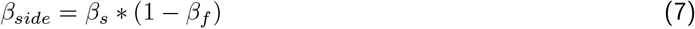

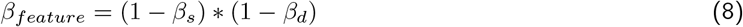

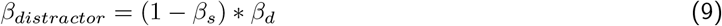

These are computed in a simple hierarchy: first *β_s_* controls whether the correct side is sampled. Second, the parameters *β_f_* and *β_d_* determine whether the patch with the feature matching the target or the distractor is sampled. We constrained *β_s_*, *β_f_*, and *β_d_* to the range [0, 1], which then also constrains *β_target_* + *β_side_* + *β_feature_* + *β_distractor_* = 1. In this way, the fit value of *β_target_* will correspond to the proportion of trials in which an observer’s response angle could be best attributed as having come from the target dot patch. *β_side_* will correspond to the proportion of trials attributed to the dot patch on the same side as the target, and similarly for *β_feature_* and *β_distractor_*.

The output of this model is then a full likelihood distribution (Fig. 4h), i.e. the probability that any given angle will be chosen as a response given the condition and stimulus (Eqn. 5).

In sum, we fit one sensitivity parameter (*d′*) and three intermediate bias parameters (*β_s_*, *β_f_*, *β_d_*) for the data set in which each observer selected by location or color (and reported motion direction) and separately for the data set in which they selected by location or motion direction (and reported color). Each model thus fit four free parameters using approximately 700 trials of data.

#### Model statistics

To compare any two variants of the models we computed their cross-validated log-likelihood ratio (i.e., the difference in total log-likelihood). We use this statistic rather than other information criterions (e.g. Akaike information criterion (Akaike, 1987)) because the cross-validation procedure already penalizes models with additional parameters for over-fitting.

To evaluate the quality of model fits for the estimation task we computed a measure of variance explained. We binned the proportion of responses at equal degree intervals (32 bins, 11.25°each) generating a distribution of response angles and compared these to the model’s predicted distribution using the formula:

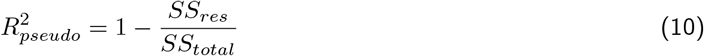

Where *SS_res_* and *SS_total_* are the unexplained variance and total variance, computed from the proportion of responses *y* and the model predictions *y′*:

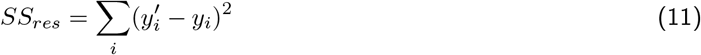

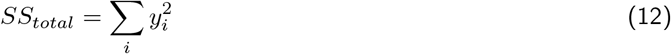

To obtain a measure of statistical significance we randomly permuted the responses made by each observer within their data and refit the models, then repeated this permutation procedure 100 times. After the subtracting the mean, the resulting distribution of log-likelihoods had a 95% CI of [−2.22, 2.54]. This matches the common suggestion that when an information criterion statistic differs by more than two it should be considered statistically significant, while a difference larger than 10 would indicate a substantial improvement in model fit. For all parameter comparisons we used permutation tests to compute confidence intervals on their differences.

#### Model recovery

To estimate the statistical power of our data set and analysis we performed a model recovery simulation. Our focus was on estimating our statistical sensitivity (i.e. true positive rate) for various effect sizes of the *d′* and *β* parameters. To estimate the sensitivity of the *d′* parameter, we set up a series of simulated data sets each consisting of 700 trials (i.e. equivalent to the data of one observer, for one task variant). These datasets were constructed by sampling response angles according to the same model used to fit the data, including the addition of motor noise (Fig. 4). We simulated a *d′* of 1.00, 1.01, 1.02, 1.03, 1.04, 1.05, 1.10, and 1.20, consistent with the range of *d′* values observed in the uncued and cued data. We set *β_target_* = 1 for these data sets. For each *d′* value we generated 200 simulated data sets and fit these with our analysis pipeline. We then compared the fit *d′* values against the distribution of values for the dataset with *d′* = 1.00. A hit was counted if the fit value for a simulated data set with *d′* > 1.00 was larger than the fit value for the data with *d′* = 1.00. We bootstrapped the comparisons 10,000 times to estimate our sensitivity and report this (markers, Fig. 6e) relative to the observed effects. The fit of a saturating exponential function (black line) which captures the simulations well is also shown.

We next set up a similar test to recover the *β* parameters. We simulated data starting from the uncued *β* values (*β_target_* = 0.50, *β_side_* = 0.35, *β_feature_* = 0.10, and *β_distractor_* = 0.05) and going up to the cued values (*β_target_* = 0.75, *β_side_* = 0.25, *β_feature_* = 0.00, and *β_distractor_* = 0.00) in 10% increments (i.e. for *β_target_*: 0% = 0.50, 10% = .525, …, up to 100%=0.75). Again we generated 200 simulated data sets for each combination of parameters, fit the model to each, and compared the parameters of each 10% increment against the fit to the 0% data, bootstrapping these 10,000 times. We calculated the proportion of simulations in which the cued *β* went in the right direction relative to the uncued *β* and report the results for the *β_target_* parameter in Figure 6f. The other *β* parameters all shared this same sensitivity curve.

## Code and data accessibility

Code and data to reproduce the analysis and figures described can be accessed with the DOI 10.17605/OSF.IO/KMBTZ.

## Acknowledgments

We acknowledge the generous support of Research to Prevent Blindness and Lions Clubs International Foundation, and the Hellman Fellows Fund to JLG as well as the UW Vision Training Grant (NEI T32EY07031) and Washington Research Foundation Postdoctoral Fellowship to DB. We thank Lynda Ichinaga for administrative support.

Current affiliation of DB: Department of Biological Structure, University of Washington, Seattle, WA 98195, USA.

## Contributions

Both the authors conceived of and designed research, interpreted results of experiments, edited and revised manuscript, and approved final version of manuscript. DB performed experiments, analyzed data, prepared figures, and drafted manuscript.

